# An open-access dataset of naturalistic viewing using simultaneous EEG-fMRI

**DOI:** 10.1101/2022.11.23.517540

**Authors:** Qawi K Telesford, Eduardo Gonzalez-Moreira, Ting Xu, Yiwen Tian, Stanley Colcombe, Jessica Cloud, Brian Edward Russ, Arnaud Falchier, Maximilian Nentwich, Jens Madsen, Lucas Parra, Charles Schroeder, Michael Milham, Alexandre Rosa Franco

## Abstract

In this work, we present a dataset that combines functional magnetic imaging (fMRI) and electroencephalography (EEG) to use as a resource for understanding human brain function in these two imaging modalities. The dataset can also be used for optimizing preprocessing methods for simultaneously collected imaging data. The dataset includes simultaneously collected recordings from 22 individuals (ages: 23-51) across various visual and naturalistic stimuli. In addition, physiological, eye tracking, electrocardiography, and cognitive and behavioral data were collected along with this neuroimaging data. Visual tasks include a flickering checkerboard collected outside and inside the MRI scanner (EEG-only) and simultaneous EEG-fMRI recordings. Simultaneous recordings include rest, the visual paradigm Inscapes, and several short video movies representing naturalistic stimuli. Raw and preprocessed data are openly available to download. We present this dataset as part of an effort to provide open-access data to increase the opportunity for discoveries and understanding of the human brain and evaluate the correlation between electrical brain activity and blood oxygen level-dependent (BOLD) signals.

## Background & Summary

Simultaneous collection of electroencephalography (EEG) and functional magnetic resonance imaging (fMRI) data is an attractive approach to imaging as it combines the high spatial resolution of fMRI with the high temporal resolution of EEG. Combining modalities allows researchers to integrate spatial and temporal information while overcoming the limitations of a single imaging modality^1,2^. Nevertheless, collecting multimodal data simultaneously requires specific expertise, and researchers must overcome various technical challenges to successfully collect data. Such challenges may limit its broader usage in the research community.

There are several technical challenges encountered when collecting imaging modalities simultaneously. With EEG, the main challenge is due to various sources of noise that impact the recorded signal. Gradient artifact is the most significant source of noise in simultaneous recordings, caused by the magnetic field gradients during fMRI acquisition, which induce current into EEG electrodes^3^. Another noise source is the ballistocardiogram (BCG) signal, which captures the ballistic forces of blood in the cardiac cycle^4,5^. The BCG artifact arises from the pulsation of arteries in the scalp that causes movement in EEG electrodes and generates voltage. The BCG artifact is more pronounced in a strong magnetic field and increases with field strength^6^. In addition to gradient and BCG artifacts, other noise sources include the MRI helium compressor^7^, eye blinks^8^, head movement, and respiratory artifacts^9^. Additionally, while collecting fMRI data, one of the main issues is patient discomfort while wearing the EEG cap in the scanner, which can cause increased head motion. Likewise, preparation time for collecting both datasets can also increase participant burden. Collecting simultaneous fMRI and EEG requires overcoming a variety of technical challenges but also needs advanced preprocessing techniques to overcome these unavoidable artifacts and produce a cleaner signal. In this paper, we detail how we addressed various technical challenges encountered when recording simultaneous EEG-fMRI including strategies to improve data quality.

For this dataset, most of the tasks performed by the participants are naturalistic viewing tasks. Naturalistic stimuli represent paradigms considered more complex and dynamic than task-based stimuli^10,11^. Naturalistic viewing provides more physiologically relevant conditions and produces closer to real-world brain responses^12–14^. Naturalistic stimuli also contain narrative structure and provide context that reflects real-life experiences^14,15^. Moreover, movies have been found to have high intersubject correlation and reliability^16,17^, holds subjects’ attention^18^, and improves compliance related to motion and wakefulness^19^. Naturalistic movies are also an ideal stimulus for multimodal data sets and may be useful in linking responses across levels^20,21^ and species^22^.

In this manuscript, we present a dataset collected at the Nathan S. Kline Institute for Psychiatric Research (NKI) in Orangeburg, NY, representing a study using simultaneously collected EEG and fMRI in healthy adults. The dataset contains multiple task conditions across two scans, including a visual task, resting state, and naturalistic stimuli. We also present quality control metrics for both modalities and describe preprocessing steps to clean up the EEG data. Lastly, we openly share these raw and processed data through the International Neuroimaging Data-Sharing Initiative (INDI) along with preprocessing code available on GitHub.

## Methods

### Participants and procedures

Simultaneous EEG-fMRI was collected in twenty-two adults (ages 23–51 years; mean age: 36.8; 50% male) recruited from the Rockland County, NY community. Participants enrolled in this study have no history of psychiatric or neurological illnesses. All imaging was collected using a 3T Siemens TrioTim equipped with a 12-channel head coil. EEG data were collected using an MR-compatible system by Brain Products consisting of the BrainCap MR with 64 channels, two 32-channel BrainAmp MR amplifiers, and a PowerPack battery. Cortical electrodes were arranged according to the international 10-20 system. Inside the scanner, eye tracking was collected in the left eye using the EyeLink 1000 Plus.

Participants attended two sessions between 2 and 354 days between scans (time between scans, mean: 38.2 days; median: 11 days); see Table 1 for the breakdown of data acquired during sessions. The scanning protocol consisted of three recording settings. The “Outside” setting was an EEG recording collected outside the MRI scanner in a non-shielded room; the “Scanner OFF” setting consisted of EEG recordings collected inside the static field of the MRI scanner while the scanner was off; the “Scanner ON” setting consisted the simultaneous EEG and fMRI recordings. All research performed was approved by NKIs Institutional Review Board. Prior to the experiment, written informed consent was obtained from all participants. Participants also provided demographic information and behavioral data, including information on their last month of sleep (Pittsburgh Sleep Study)^23^, the amount of sleep they had the previous night, and their caffeine intake before the scan session.

**Table 1.** Simultaneous EEG-fMRI experimental design. EEG and fMRI were collected simultaneously across two scan sessions.

| Session | Recording Location | Scan | Run Length (s) |
| --- | --- | --- | --- |
| Day 1 | Outside | Checkerboard (12Hz) | 200 |
|  | Scanner OFF | Checkerboard (12Hz) | 200 |
|  | Scanner ON | Checkerboard (12Hz) | 200 |
|  |  | Rest | 600 |
|  |  | Inscapes | 600 |
|  |  | The Present [Run 1] | 258 |
|  |  | PEER | 90 |
|  |  | Monkey 1 [Run 1] | 300 |
|  |  | Despicable Me (English) [Run 1] | 600 |
|  |  | MPRAGE | 600 |
|  |  | The Present [Run 2] | 258 |
|  |  | Monkey 1 [Run 2] | 300 |
|  |  | Despicable Me (English) [Run 2] | 600 |
| Day 2 | Outside | Checkerboard (12Hz) | 200 |
|  | Scanner OFF | Checkerboard (12Hz) | 200 |
|  | Scanner ON | Checkerboard (12Hz) | 200 |
|  |  | Rest | 600 |
|  |  | Inscapes | 600 |
|  |  | Monkey 2 [Run 1] | 300 |
|  |  | PEER | 90 |
|  |  | Despicable Me (Hungarian) [Run 1] | 600 |
|  |  | Monkey 5 [Run 1] | 300 |
|  |  | MPRAGE | 600 |
|  |  | Monkey 2 [Run 2] | 300 |
|  |  | Despicable Me (Hungarian) [Run 2] | 600 |
|  |  | Monkey 5 [Run 2] | 300 |

### EEG acquisition

The entire procedure of data collection takes approximately three hours. About 45 minutes are spent preparing a participant for EEG. Preparation begins with measuring the participant’s head to determine cap size. An anterior to posterior measurement is also taken across the top of the head, from the nasion to the inion. A mark is made at the center of the forehead 10% of the measured nasion-inion distance. A proper sized cap is placed on the participant’s head with the Fpz electrode centered at the mark on the forehead. Once the subject is fitted with the EEG cap, electrodes are filled with electrolyte gel. In this study, EEG is collected using a customized cap to record 61 cortical channels, two electrooculogram (EOG) channels placed above (channel 64) and below the left eye (channel 63), and one electrocardiography (ECG) channel (channel 32) placed on the back. In addition, the cap also contains a reference and ground electrode. Electrodes were filled using V19 Abralyt HiCl electrode gel. Electrode impedance was recorded before every recorded run; to ensure good data quality, electrode impedance was kept below 20kOhm. EEG was recorded using BrainVision Recorder at a sampling rate of 5 kHz. After cap preparation, participants completed a single run of the flickering checkerboard experiment in the “Outside” scan condition. After the Outside scan, a 3D scan of the participant’s head is collected to digitize the position of EEG electrodes. 3D scans were collected using a portable scanner, the Occipital Structure Sensor (Occipital Inc, Boulder CO), and iPad Mini 4 (Apple, Cupertino, CA). Due to protected health information (PHI) restrictions, 3D scans will not be available in the data release; however, location files containing the positions of the electrodes will be provided with this release.

### Simultaneous EEG-fMRI recording in the MRI scanner

After 3D digitization, participants enter the MRI scanner and are placed on the scanner bed in the supine position. Cushions are placed around the head to provide stabilization and minimize head motion during scans. At this stage, participants are provided with MR-safe goggles if they have any visual impairment that requires glasses. Participants were also fitted with a respiratory transducer belt to monitor breathing, which was recorded using BIOPAC MP150 (BIOPAC Systems, Inc., Goleta, CA).

Once positioned in the scanner, the participant’s head is enclosed in a 12-channel 3T head matrix MR coil. The cable bundle from the EEG cap is routed through the front of the head coil and fixed at the top of the head coil using medical tape. The cable bundles are connected to the amplifiers and battery pack. During EEG data acquisition, the software is synchronized to the master clock of the MRI scanner. During recordings, EEG data was continuously collected along with task onset triggers, and volume triggers were recorded at the beginning of each TR. For specifics on the equipment and connections for simultaneous recordings, see Figure 1 and Table 2.

**Figure 1.**
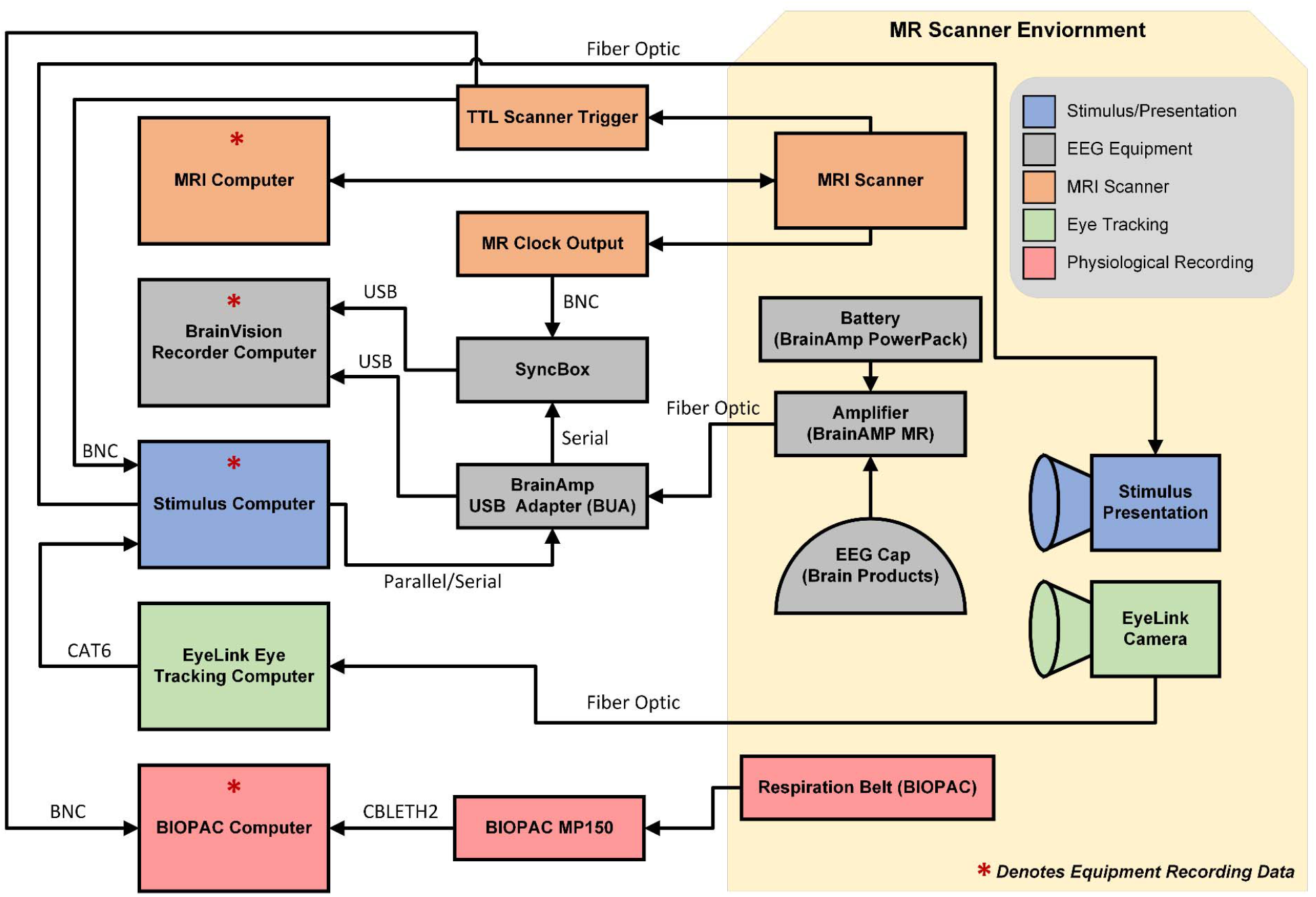
Schematic of EEG-fMRI setup.

### Eye tracking acquisition

For recordings collected inside the scanner, eye position and pupil dilation were recorded using an infrared-based eye tracker (EyeLink 1000 Plus, SR Research Ltd., Ontario, Canada; http://www.sr-research.com) at a sampling rate of 1000Hz. The eye tracker was calibrated using a 9-point grid before recordings in the MRI scanner. Participants were asked to direct their gaze at dots presented on the grid. Calibration was followed by a validation step until the error between the two measurements was less than 1° ^24^.

### MRI data acquisition

MRI data were acquired using a 12-channel head coil on 3.0T Siemens TIM Trio. MPRAGE structural T1w images were acquired with the following parameters: TR = 2500 ms; TI = 1200 ms; TE = 2.5 ms; slices = 192; matrix size = 256×256; voxel size = 1 mm^3^ isotropic; flip angle = 8°; partial Fourier off; pixel bandwidth = 190 Hz/Px. All BOLD fMRI sequences were acquired with these parameters: TR = 2100 ms; TE = 24.6 ms; slices = 38; matrix size = 64×64; voxel size = 3.469×3.469×3.330 mm. The run length for each task is listed in Table 1.

### Task data/stimuli description

In this section, we describe the data generated for this study focused on collecting simultaneous EEG and fMRI. This dataset consists of a task, naturalistic stimuli, and resting state data. In addition to the simultaneously collected data, task data was collected outside the MRI scanner and inside the scanner environment with the scanner off. The collection of this data enables us to assess the impact of changes in the scanning environment on the EEG recordings. EEG-fMRI data were collected across two scan sessions; structural data was collected during the middle of the scan (see Table 1 for details). Code for presenting task stimuli and naturalistic stimuli, along with code to preprocess EEG and fMRI imaging data is available on GitHub (https://github.com/NathanKlineInstitute/NATVIEW_EEGFMRI).

#### Checkerboard stimulus

The use of flickering visual stimuli has been used to investigate the visual system in EEG^25,26^ and fMRI^27,28^. A high-contrast flickering checkerboard was used to stimulate the primary visual cortical regions. Participants were shown a flickering radial checkerboard at a frequency of 12Hz in 20-second trials following a 20-second rest period across five repetitions. The checkerboard stimulus was presented outside the scanner after cap placement, inside the scanner with the scanner off, and inside the scanner during a simultaneous EEG and fMRI. These three recordings were collected to measure the impact of the MRI scanning sequence on the recorded EEG.

#### Rest

The participant is presented with a white fixation cross in the center of a black screen and instructed to rest with eyes open. Participants had one rest scan per session, and each scan had a duration of 10 minutes.

#### Inscapes

Inscapes is a computer-generated animation featuring abstract 3D shapes and moves in slow continuous transitions. The video was originally developed as a 7-minute video for children to watch during brain scans as a means to provide stimulation to keep them engaged while minimizing some cognitive processing that may be engaged^19^. Participants were presented with an extended version of Inscapes that was 10 minutes long. Similar to the rest scan, the Inscapes video was viewed once per scan session.

#### Predictive eye estimation regression (PEER) calibration

Predictive eye estimation regression (PEER) is an imaging-based calibration scan used to estimate the direction of eye gaze^29^ and here can be used as a complement to the optical eye tracking of the Eyelink 1000. Participants were told to direct their gaze at dots that appeared at predefined points on the screen. The PEER method estimates eye gaze from the collected scan using support vector regression (SVR). The algorithm estimates the direction of eye gaze during each repetition (TR) in the fMRI time series.

#### Naturalistic stimuli (Movies)

Participants viewed three movies twice in one scan session. Videos varied between 258s and 600s. Naturalistic stimuli included “The Present” [4m18s] (uploaded to YouTube 7 Feb 2016)^30^, two 10-minute clips from “Despicable Me” [clips taken from Russian Blu-Ray with exact times 1:02:09-1:12:09 (English) and 0:12:12-0:22:12 (Hungarian)]^31^, and three 5-minute monkey videos^32^. The monkey videos are part of a database with multiple videos^33^; for this study, Monkey 1, Monkey 2, and Monkey 5 represent the first, second, and fifth video in the database, respectively. The videos used in this study are available to download in the GitHub repository (https://github.com/NathanKlineInstitute/NATVIEW_EEGFMRI/tree/main/stimulus).

### Limitations

The fMRI imaging in this dataset was collected using a 12-channel head coil on a 3T scanner platform. While we possess the ability to scan at 3T with a 32-channel head coil, the head coil design does not allow for the EEG cap cable bundle to be routed perpendicularly from the top of the head. Moreover, the design of the head coil does not permit alternate routing of the cable bundle because it will block the participant’s eyes or the cable length is too short to reach the amplifiers. In this study, the cable bundle is routed through the front and then taped to the top of the head coil before it is connected to the amplifier. Using a 12-channel head coil limited the sequences that could be used to collect fMRI, including the use of multiband sequences. Another limiting factor is that imaging sequences with faster TRs could not be used while collecting data simultaneously. The main issue is a safety concern regarding radiofrequency (RF) power deposition that causes heating in the EEG leads and electrodes during a sequence^34^. In this study, we used a TR of 2100ms to ensure participants were not at risk of discomfort or burns due to the heating of electrodes.

### EEG preprocessing

We developed an automated pipeline to preprocess all EEG data collected outside and inside the MRI scanner. The preprocessing methods used on the EEG data depended on where data were acquired. All EEG data were preprocessed using EEGLAB and associated plugins^35^. For data collected inside the scanner, data were preprocessed using the FMRIB plug-in for EEGLAB, provided by the University of Oxford Centre for Functional MRI of the Brain (FMRIB)^36,37^.

For data collected in the Outside setting, the following preprocessing steps were used: (i) bandpass filter using a Hamming windowed sinc FIR filter between 0.5Hz and 70Hz; (ii) reference electrodes using average reference, excluding the ECG channel, EOG channels, and electrodes excluded during the EEG quality control process.

For data collected in the Scanner OFF setting, the initial preprocessing steps used in the Outside setting were used. In addition, pulse artifact detection and removal were used due to the increased contamination of the signal caused by the participant’s heartbeat^3^: (i) QRS/heartbeat detection using the ECG channel; (ii) pulse artifact/BCG removal using template subtraction based on the median artifact; (iii) bandpass filter using a Hamming windowed sinc FIR filter between 0.5Hz and 70Hz; (iv) reference electrodes using average reference, excluding the ECG channel and EOG channels.

For data collected in the Scanner ON setting, the initial preprocessing steps used in the Scanner OFF setting were used. In addition, gradient artifact removal was used to remove the contamination of the EEG signal caused by the changing gradients from the fMRI pulse sequence^36^: (i) gradient artifact removal; (ii) QRS/heartbeat detection using the ECG channel; (iii) pulse artifact/BCG removal using template subtraction based on the median artifact; (iv) bandpass filter using a Hamming windowed sinc FIR filter between 0.5Hz and 70Hz; (v) reference electrodes using average reference, excluding the ECG channel and EOG channels.

#### Gradient artifact removal

The gradient artifact is the most significant noise source in simultaneous EEG-fMRI data, measuring more than 400 times larger than the lowest amplitude EEG events^3^. The FMRIB plugin uses the FASTR method to remove gradient artifacts^37^. The method requires recording the scanner trigger at the start of each TR. An average template is computed from the detected TRs and subtracted from the raw EEG data. Following this process, the data is corrected further using principal component analysis (PCA) to reduce residual artifacts. Residual artifacts are further reduced using adaptive noise cancellation^3^.

#### Pulse artifact/BCG removal

ECG data collected inside the MRI scanner has a pronounced T wave that increases as the field strength increases^38^. The FMRIB plugin identifies these QRS events using an algorithm that detects events, aligns them, and corrects for false positives and negatives^39,40^. A median signal is computed from the events to create an artifact template, which is subsequently subtracted from the data.

### MRI preprocessing

We used the Connectome Computation System (CCS) to preprocess the MRI/fMRI data^41^. For the anatomic data, we performed skull stripping using a combination of Brain Extraction Toolbox (BET) and Freesurfer. Data was then segmented (Freesurfer) and registered to a template space (MNI152 2006) using FLIRT and MCFLIRT^42,43^. The runs for the fMRI data were all preprocessed equally. Initially, the first five volumes are discarded, then the data is despiked and slice time and motion corrected. The functional data is skull striped with 3dAutomask, refined using the structural data, and registered to the anatomical images using boundary-based registration based on N4 using Freesurfer. Nuisance correction is done using the Friston 24 motion parameters, average CSF, and WM signals, with/without global signal regression (GSR). The data is also processed with/without temporal filtering (0.01-0.1Hz) and with/without a 6mm FWHM spatial filter. Time series were extracted from 400 ROIS (Schaefer 400 atlas) to be further processed^44^.

### Data quality control

#### EEG

To assess the quality of the EEG data in this study, we followed the quality control pipeline similar to that described in Delorme *et al*^45^. This approach yields three metrics related to data quality: (i) percent of “good” channels; (ii) percent of “good” trials; and (iii) number of independent components (ICs) related to brain source activity. From this pipeline, “good” channels are defined as those that remain after completing the related preprocessing steps: removal of channels with more than five seconds of non-activity, with signal greater than four standard deviations due to high-frequency noise, or Pearson’s correlation coefficient less than 0.7 with nearby channels. Accordingly, “good” trials are related to the data periods that are not contaminated by artifacts such as body movement. In this study, we removed data segments with a variance higher than twenty times the variance of the calibration data. Finally, independent component analysis (ICA) was computed using the RunICA plugin for the EEGLAB toolbox to produce ICs^46^. Afterward, the ICLabel plugin was used to identify ICs belonging to brain source activity^47^. The resulting metric calculates the percentage of ICs associated with brain source activity divided by the total number of ICs found from ICA.

#### MRI

Temporal measures of fMRI data include median and median framewise displacement^43^, root mean square of the temporal change (DVARS), and temporal signal-to-noise ratio (tSNR).

### Statistical analysis

Permutation testing offers a robust framework for statistical significance assessment in EEG analysis. Multiple permutation testing was performed on the flickering checkerboard task to identify differences between the two task conditions: the rest and flickering checkerboard blocks. This analysis is used to find statistical relevance that identifies differences underlying the EEG data. One advantage of multiple permutation testing is that it does not require the same number of trials for each condition. In our permutation testing, trials were shuffled into two new groups, followed by the calculation of a paired t-test. Finally, pixel-based multiple comparisons correction was applied to reduce the familywise error rate^48^.

## Data Records

### Open-access data

There are various neuroimaging databases providing open-access datasets, such as OpenNeuro (https://openneuro.org), DataDryad (https://datadryad.org), and Zenodo (https://zenodo.org). Across the available datasets (see Table 3), EEG-fMRI data exists for sleep, tasks, and resting state data^1,49–62^. While there are some simultaneous datasets available, to our knowledge, no open-access datasets are available with naturalistic viewing data. This dataset is more comprehensive, and contains resting state, a visual task paradigm, and naturalistic video stimuli. In our data release, we provide raw and preprocessed data. In addition, physiological data collected concurrently is also provided, including eye tracking, electrooculography, electrocardiography, and respiration rate. For a description of the data collected in this study, see Table 2.

### Data privacy

All imaging data in this release has been de-identified, removing any personal identifying information (as defined by the Health Insurance Portability and Accountability) from data files, including facial features. Facial features from the T1w MRIs were removed using the “mri_deface” software package developed by Bischoff-Grethe *et al*.^63^. Data and code are shared under the CC BY-SA 4.0 license.

### Data access

All data can be accessed through the 1,000 Functional Connectomes Project and its International Neuroimaging Data-sharing Initiative (FCP/INDI) based at https://fcon_1000.projects.nitrc.org/indi/retro/nat_view.html. This website provides directions for users to directly download all the imaging data from an Amazon Simple Storage Service (S3) bucket. Raw data is stored in the Brain Imaging Data Structure (BIDS) format^64^, which is an increasingly popular approach to describing imaging data in a standard format. Preprocessed data is also provided through the website.

**Table 2.** Equipment used for data collection and stimuli presentation in EEG-fMRI study.

| **Modality** | **Equipment** | **Sampling Rate** | **Additional Information** |
| --- | --- | --- | --- |
| EEG/EOG/ECG | Brain Products BrainCap MR with Multitrodes | 5000Hz | 64 channels<br>• EEG: 61 cortical electrodes<br>• EOG: 2 channels<br>◦ Placed below and above left eye<br>◦ EOGL (Lower: channel 63)<br>◦ EOGU (Upper: channel 64)<br>• ECG: 1 channel<br>◦ placed on participant's back below left shoulder blade (channel 32) |
| fMRI | Siemens 3.0T TIM Trio | 0.476Hz<br>(TR = 2100ms) | 12-channel head coil<br>• TR = 2100ms<br>• TE = 24.6ms<br>• slices = 38<br>• matrix size = 64x64<br>• voxel size = 3.469x3.469x3.330 mm |
| Eye Tracking | EyeLink 1000 Eye Tracker | 250Hz or 1000Hz | Left eye |
| Respiration | BIOPAC MP150 | 62.5Hz | Placed around abdomen/chest |
| Stimulus Presentation | NEC LT220 DLP Projector | N/A | Video displayed at<br>Display Resolution: 1024x768<br>Refresh Rate: 60Hz |
| 3D scans | Occipital Structure Sensor | N/A | 3D scanner paired with Apple iPad Mini 4 |

**Table 3.** Open-access simultaneous EEG-fMRI data sets. Simultaneous EEG-fMRI data sets are available from various online repositories on OpenNeuro, DataDryad and Zenodo for sleep, task, and resting state data.

| Authors | Task | Participants | Scanner | Functional | Head Coil | EEG Cap | Physiological Measures | Notes |
| --- | --- | --- | --- | --- | --- | --- | --- | --- |
| Robertson et al., 2015<sup>49,50</sup> | multi-source interference task (MSIT) | 18 healthy (17 right-handed, 1 left-handed; 10 females, 8 males) | 3T Siemens Verio (Siemens, Erlangen, Germany) | EPI [TR: 3000ms; TE: 30ms; flip angle: 90°; matrix size: 64×64; voxel size: 3.2 × 3.2 × 3.2 mm; 30 slices, butting] | 12-channel phased array head coil | Neuroscan (64 electrodes) | ECG, EOG | - |
| Zumer et al., 2015<sup>51,52</sup> | one-back visuospatial working memory task | 16 healthy participants (15 females) | 3T Siemens TIM Trio (Siemens, Erlangen, Germany) | EPI [TR: 2000ms; TE: 6.9, 16.2, 25, 35, 44 ms; flip angle: 80°; matrix size: 64×64; voxel size: 3.5 × 3.5 × 3 mm; 39 slices; acceleration factor: 3] | 32-channel head coil | BrainProducts (64 electrodes) | ECG, infrared eye tracker | Polhemus FASTRAK device was used to record the exact location of each EEG electrode on the participant's head relative to three fiducial points |
| Roy et al., 2017<sup>53,54</sup> | 4m42s (5 consecutive 42s trials followed by 12s rest) [2538s (9) - 4794s (14)] | 20 healthy | 3T Siemens Skyra (Siemens, Erlangen, Germany) | EPI [TR: 2200ms; TE: 30ms; flip angle: 90°; voxel size: 3 × 3 × 3 mm; 36 slices; fat suppression on] | 16 channel receive only head coil | BrainProducts (64 electrodes) | ECG | 12 preprocessed EEG scan files + 12 preprocessed fMRI scan folders |
| Gherman and Philiastides, 2018<sup>55,56</sup> | perceptual decision task (random dot kinematogram) | 30 healthy (24 used in study) | 3T Siemens TIM Trio (Siemens, Erlangen, Germany) | EPI [TR: 2000ms; TE: 30ms; flip angle: 80°; FOV: 210mm; matrix size: 70×70; voxel size: 3 × 3 × 3 mm; 32 slices, interleaved; 0.3 mm slice gap] | 12-channel head coil | BrainProducts (64 electrodes) | ECG | - |
| He et al., 2018<sup>57,58</sup> | content judgment task (2 runs, 425 volumes) | 20 participants (12 females) | 3T Siemens TIM Trio (Siemens, Erlangen, Germany) | EPI [TR: 2000ms; TE: 30ms; flip angle: 90°; FOV: 230mm; matrix size: 64×64; voxel size: 3.59 × 3.59 × 4 mm; 30 slices; 0.36mm slice gap] | Unspecified | Brain Products (32 electrodes) | ECG | - |
| Lioi et al., 2019<sup>59</sup> | 6m40s neurofeedback, 5m20s neurofeedback (5 runs) | 30 right-handed NF-naïve healthy | 3T Verio MRI [VB17] (Siemens, Erlangen, Germany) | EPI [TR: 2000ms; TE: 23ms; FOV: 210mm; voxel size: 2 × 2 × 4 mm; 32 slices; no slice gap] | 12-channel receiver head coil | - | - | Only 27 of them signed the consent to publish their anonymized data, and were therefore included in the public datasets |
| Mele et al., 2019<sup>†</sup> | attention task (380 days; range 239–560 days) | 5 healthy | 3T Philips Achieva (Philips Medical Systems, Best, Netherlands) | EPI [TR: 2000ms; TE: 30ms; flip angle: 90°; matrix size: 80×80; voxel size: 2.75 × 2.75 × 3.75 mm; 36 slices, acquired continuously in ascending order; no slice gap] | 32-channel head volume coil | WaveGuard (64 electrodes) | ECG, EOG | - |
| Wirsich et al. 2020<sup>60,61</sup> | resting state | 16 healthy | 1.5T Siemens Avanto (Siemens, Erlangen, Germany) | EPI [TR: 2160ms; TE: 30ms; flip angle: 75°; FOV: 210mm; voxel size: 3.3 × 3.3 × 4 mm; 30 slices; 1mm slice gap] | Unspecified | BrainProducts (64 electrodes) | ECG | Only 1.5T available, other data available upon request |
|  | resting state | 26 healthy | 3T Siemens TIM Trio (Siemens, Erlangen, Germany) | EPI [TR: 2000ms; TE: 50ms; flip angle: 78°; FOV: 192mm; voxel size: 3 × 3 × 3 mm; 40 slices] | Unspecified | BrainProducts (64 electrodes) | ECG, EOG | - |
|  | resting state | 21 healthy | 3T Siemens TIM Trio, 3T Siemens Prisma (Siemens, Erlangen, Germany) | EPI [TR: 1990ms; TE: 30ms; flip angle: 90°; FOV: 192mm; voxel size: 3 × 3 × 3.75 mm; 32 slices] | Unspecified | Electrical Geodesic (258 electrodes) | 2 ECG | - |
|  | resting state | 9 healthy | 7T Siemens Magnetom [head MR-scanner] (Siemens, Erlangen, Germany) | EPI [TR: 1000ms; TE: 25ms; flip angle: 54°; FOV: 220mm; voxel size: 2.2 × 2.2 × 2.2 mm; 69 slices; acceleration factor: 2; simultaneous multi-slice: 3] | Unspecified | BrainProducts (64 electrodes) | ECG, 4 modified motion | - |
| Gu et al., 2022<sup>62</sup> | 10m resting state (2 runs); 15m sleep (varied) | 33 healthy | 3T Siemens Prisma Fit (Siemens, Erlangen, Germany) | EPI [TR: 2100ms; TE: 25ms; FOV: 240mm; voxel size: 3 × 3 × 4 mm; 35 slices] | 20-channel receive-array | Brain Products (32 electrodes) | ECG, EOG | Structural and functional MRI, Raw EEG |

## Technical Validation

### EEG data validation and quality

Figure 2 shows the comparison of EEG during the checkerboard experiment across the three scan settings. Figure 2A shows the signal power over 2s epochs during the checkerboard and rest task averaged across subjects at the Oz electrode. Comparing the checkerboard versus rest condition, the checkerboard shows peaks at 12Hz and 24Hz, representing the driving frequency of the checkerboard and its harmonic, respectively. When moving into the scanner environment, the power of the checkerboard is reduced. Nonetheless, peaks at 12Hz and 24Hz are still visible in the Scanner OFF and Scanner ON settings. The Scanner ON setting also contains a dip at 18Hz in both rest and checkerboard conditions. This dip denotes residual artifacts that are left behind in the signal after the gradient artifact removal step. When looking at the frequency over the duration of the 2s epoch, the difference between the rest and checkerboard condition is evident with the 12Hz driving frequency and the 24Hz harmonic appearing across the entire epoch (Figure 2B). As shown in Figure 2C, these differences are statistically significant for the three settings. While these plots focus on the Oz electrode, the signal power also extends to other electrodes in the occipital region (Figure 2D).

**Figure 2.**
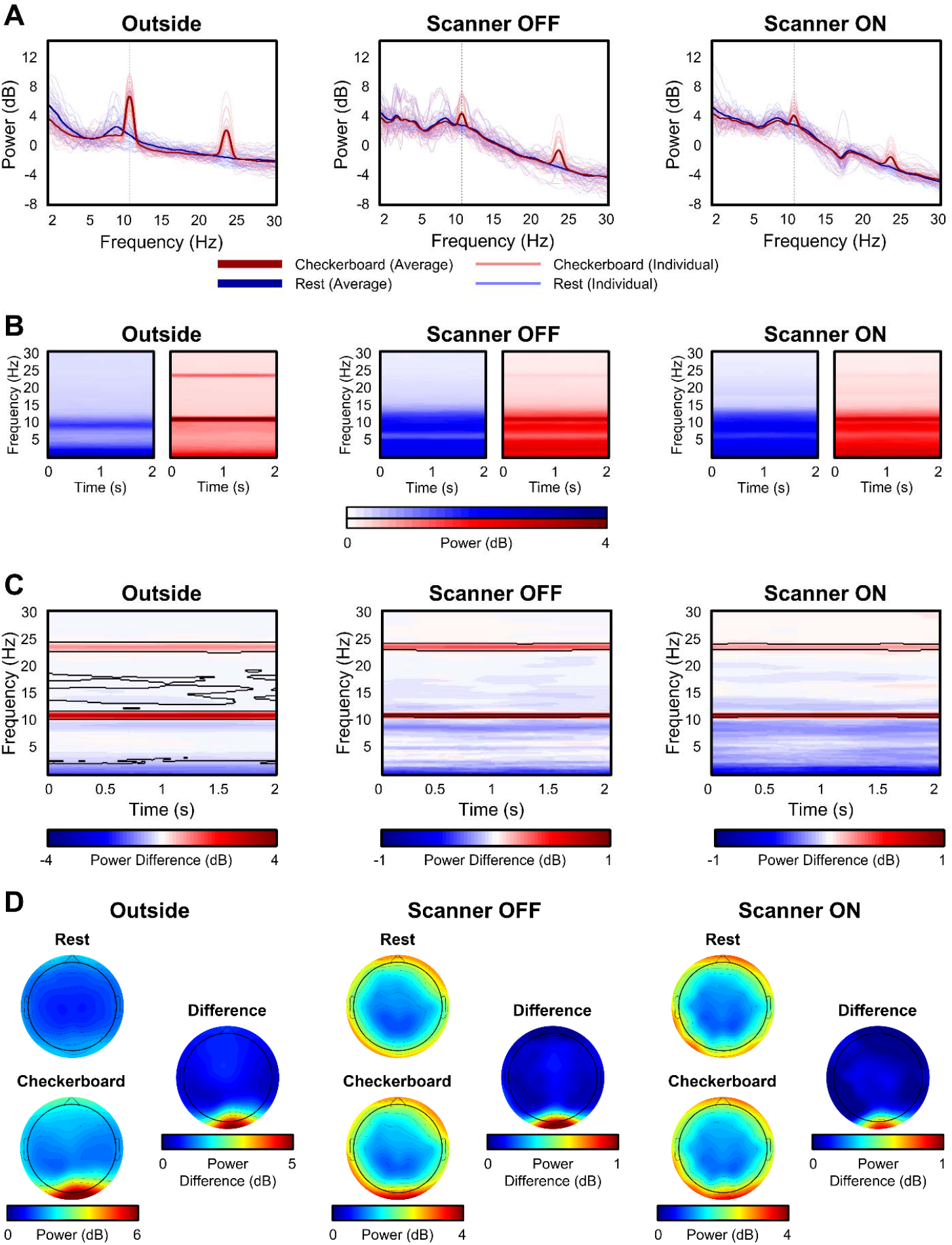
EEG data validation. A) Comparison of power spectrum for participants across scanning conditions: Outside, Inside the scanner with the Scanner OFF, and Scanner ON; B) Time-frequency comparison of checkerboard and rest blocks over 2s epochs across conditions; C) Permutation subtraction of Checkerboard and Rest. Statistically significant region are outlined by black lines. It is notable that the frequency range around 12 and 24Hz are statistically significant across all conditions; D) Topographic plots comparing Rest and Checkerboard condition, and their difference, across three conditions.

Three quality assessment metrics were computed for each raw EEG dataset: percent of “good” channels, percent of “good” trials, and the number of independent components (ICs) related to brain source activity as a percentage of the total number of ICs. As shown in Figure 3, data quality was high across all subjects for the percentage of good channels and trials for the checkerboard task. Although the data quality was highest in the Outside setting, high percentages were found for channels and trials in the Scanner OFF and Scanner ON settings. Similarly, as seen in Figure 4, the percentage of good channels and trials was high across tasks, denoting the stability of data quality during the scan session. The percent of putative brain sources based on ICs classification was lower for the Scanner ON setting compared to the other two settings. Due to the increase in noise sources in the Scanner OFF (e.g., pulse artifact) and Scanner ON (e.g., gradient artifact) settings, the percentage of ICs related to brain sources is expected to decrease. As shown in Figure 4, the quality of the EEG data is stable across scan settings.

**Figure 3.**
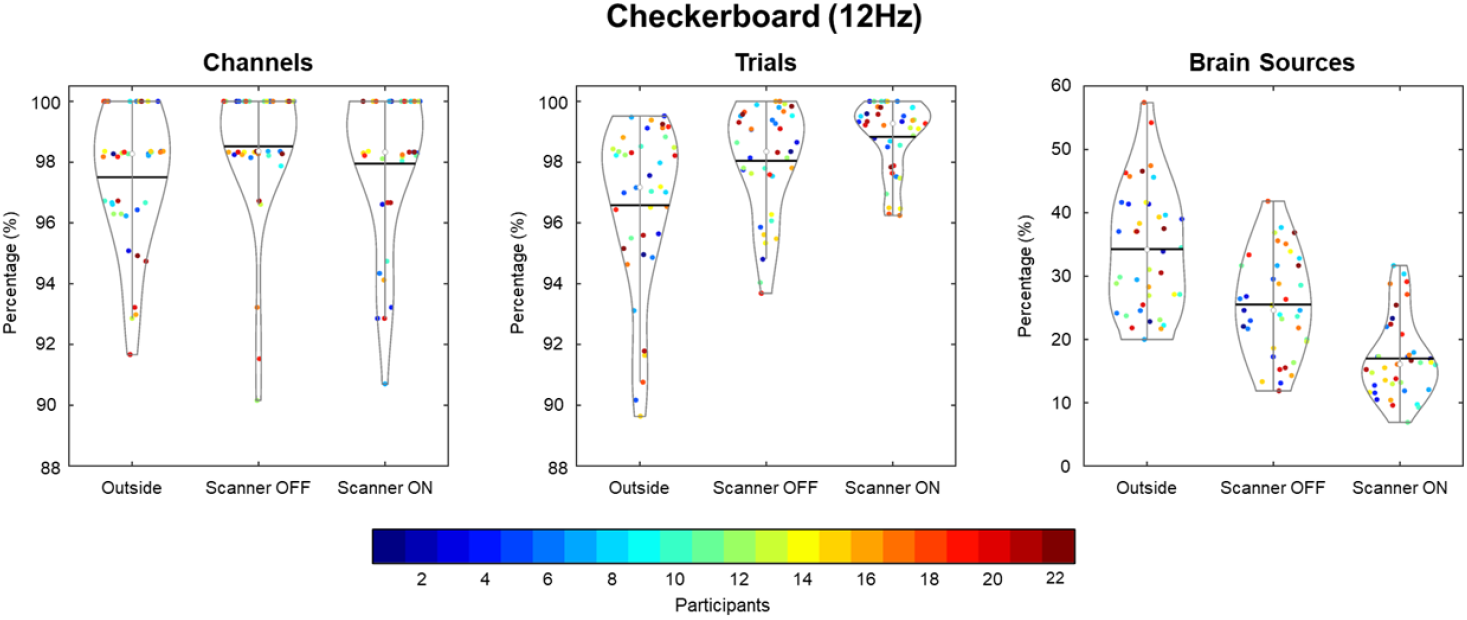
EEG data quality comparison for flickering checkerboard task across participants. Three quality metrics (percentage of good channels, percentage of good trials and percentage of ICA brain sources) were assessed across scan conditions. For channels and trials, the data quality was high across all scan conditions, denoting high quality of EEG data.

**Figure 4.**
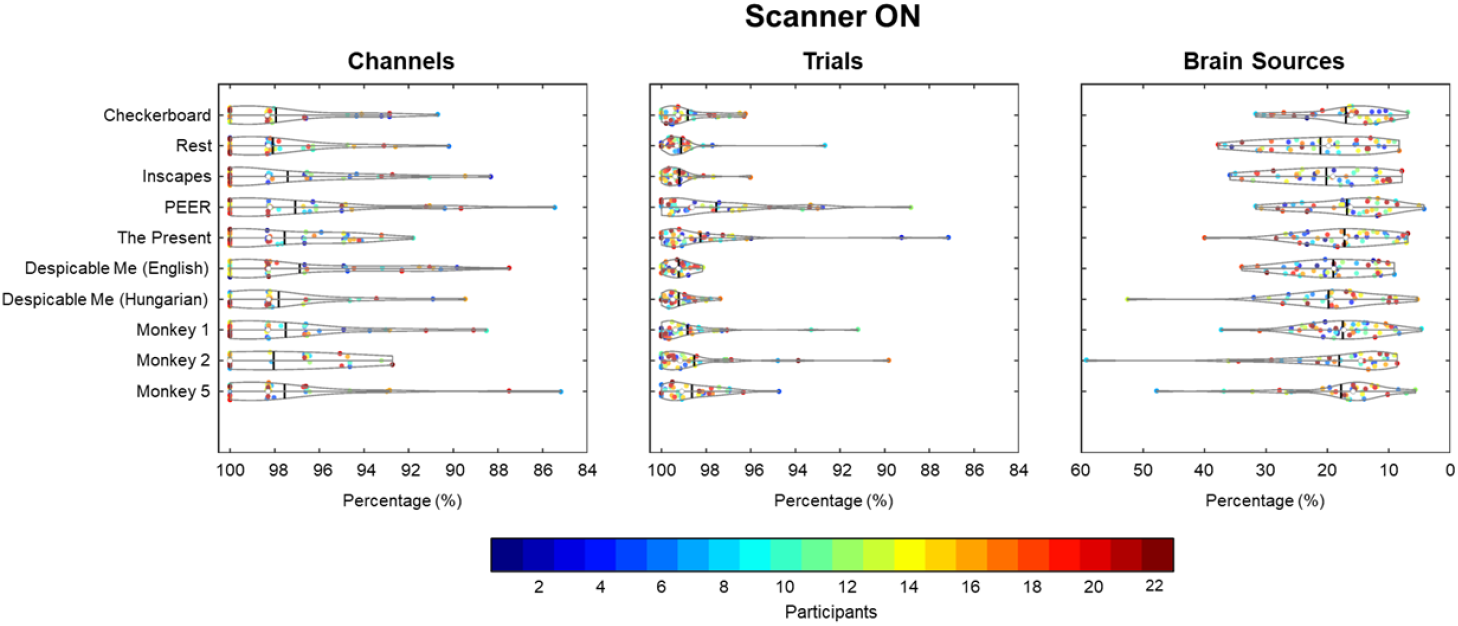
EEG data quality comparison inside scanner. Three quality metrics (percentage of good channels, percentage of good trials and percentage of ICA brain sources) were assessed across scan conditions. For channels and trials, the data quality was high across all scan conditions, denoting maintenance of high quality of EEG data across the imaging session.

### fMRI data quality

To assess the quality of fMRI data, median framewise displacement (FD) was measured for all scans. As shown in Figure 5, the median FD was for every fMRI scan; scans with a value above 0.2 were considered high motion. To determine if there was an ordering effect, scan sessions were color coded to determine if participants moved more earlier or later in the scan. Most subject data were below the 0.2 threshold (93% of scans), and there was no pattern of ordering across participants.

**Figure 5.**
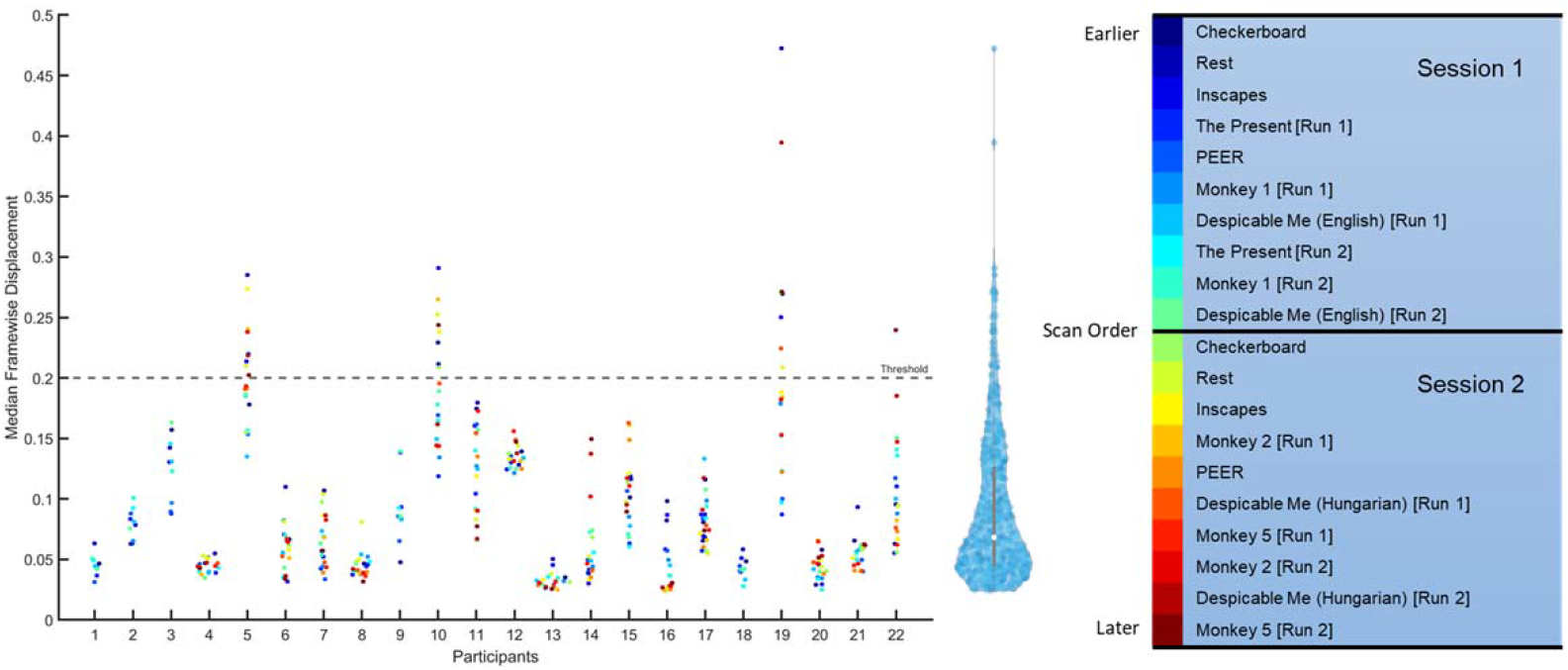
Median framewise displacement for fMRI data across the 22 participants in the study. The median framewise displacement was measured for each scan across both sessions and plotted for each participant. Scans with a value above 0.2 were considered high motion, indicated as points about the dotted threshold line. To the right of the plot, is the distribution of all scans. As seen in the distribution, participant compliance was good and most scans had low motion as measured by median FD.

For the checkerboard experiment, we looked at the correlation between ROIs within and between subjects and across scans (Figure 6). The distributions for within-scan and within subjects showed a broader distribution of values, with higher correlations for within-scan and within subject distributions.

**Figure 6.**
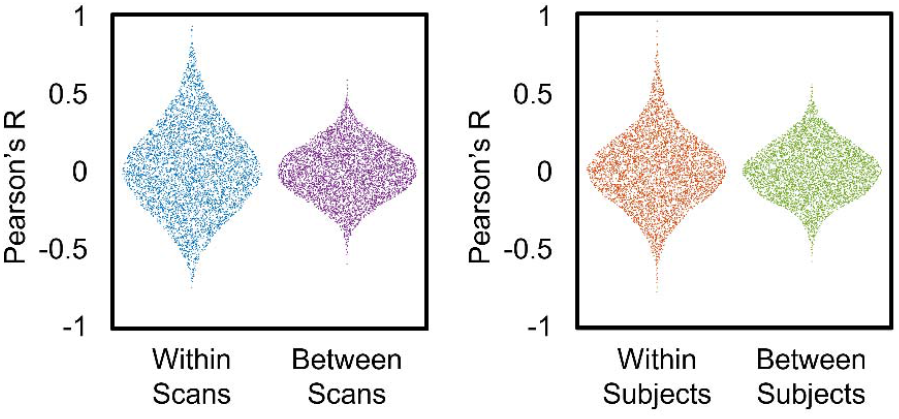
Distributions of correlation coefficients comparing within scans and between scans from the same participant, and within subject and between subject scans across all participants. When viewing a single participant, the distribution of correlation coefficients is broader (i.e., a longer tailed distribution) within scan compared to correlations between scans, reflecting stronger intrascan correlation coefficients. Similarly, the within subject correlation coefficients were stronger within participant compared to correlations between subjects.

### Multimodal quality data correlations

Values for EEG and MRI data were compared within and between modalities across several quality metrics: mean FD (framewise displacement), median FD, DVARS (temporal derivative of time courses), and tSNR (temporal signal-to-noise ratio) for MRI; channels, trials, and brain sources for EEG (Figure 7). Using Spearman’s *ρ* between each modality, shows a strong positive correlation between mean and median FD, and a strong negative correlation between FD measures and tSNR. A weak correlation was found between DVARS and tSNR, but no association was found with DVARS and other measures. For EEG measures, there was no correlation between the different quality measures. Moreover, there was no correlation between quality measures between imaging modalities..

**Figure 7.**
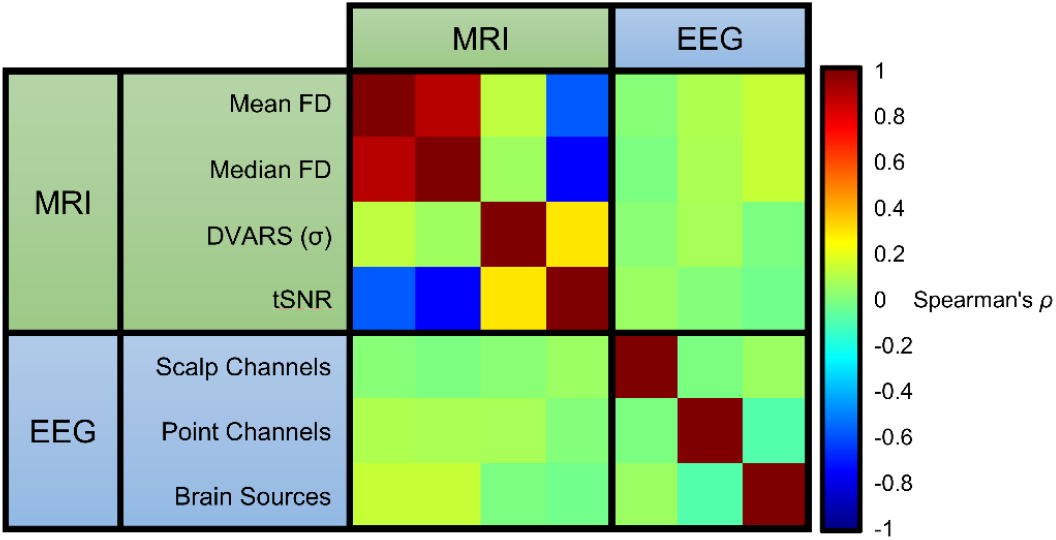
Correlation between MRI and EEG data. As expected, there is strong positive correlation between measures of mean and median FD; likewise, there is a strong negative correlation between measures of FD and tSNR. In contrast, there appears no association among EEG measures within modality and with fMRI quality measures.

### Multimodal data integration

As a test for multimodal data integration, we evaluated whether we could use the EEG signal to predict the hemodynamic response in the fMRI data. Specifically, after preprocessing, the EEG signal from Oz signal was averaged across participants for the checkerboard experiment. This signal was then bandpass filtered (20th order IIR filter between 11Hz and 13Hz), modulated, and convolved with an ideal hemodynamic response function (using a gamma variate function). This signal was used as a regressor for each participant to map out the BOLD activity. A one-sample t-test was performed to calculate a group activity map (Figure 8A). For comparison purposes, a regressor based on an ideal block design convoluted a gamma variate function was also calculated to look at the group-level activity (Figure 8B). Activity maps shown in Figure 8 indicate that both approaches generate a similar level of activity in the occipital lobe.

**Figure 8.**
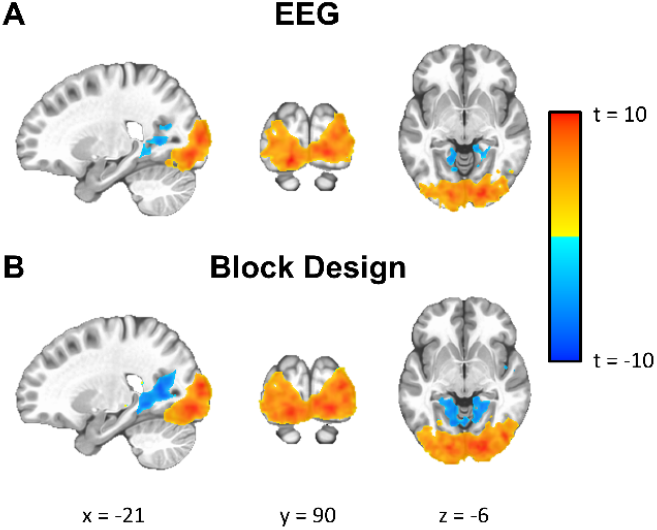
Checkerboard task group activation map either using the A) group average EEG signal from Oz as a regressor or B) a block design to generate the hemodynamic response function.

### Technical challenges

Collecting EEG and fMRI simultaneously requires several methodological considerations. While EEG and fMRI have a long-established history, collecting EEG inside the MRI scanner is challenging for several technical reasons. The main problem encountered when collecting a functional recording is the generation of artifacts from various sources. The main artifact arises from the gradient artifact generated during echo-planar imaging (EPI), which induces changes in the magnetic field^65^. Another source of noise arises from the scanner environment. While not a problem in all scanners, vibrations from the helium compressor in Siemens Trio and Verio scanners introduce artifacts into the EEG signal^66^; these vibrations induce non-stationary artifacts that contaminate the EEG signal. Yet another source of noise is caused by the pulsation of arteries in the scalp that cause movement in EEG electrodes and generation voltage. The ballistocardiogram (BCG) signal captures the ballistic forces of blood in the cardiac cycle^4,5^ and becomes more pronounced as the magnetic field strength increases^6^. In addition to a more pronounced signal, the ECG signal can impact data collection and preprocessing. In some cases, the pronounced ECG signal leads to saturation of the signal during the MRI scan sequence. Consequently, this saturation causes signal clipping that impedes QRS detection and pulse artifact removal methods during preprocessing. In this data release, there are occasions of signal clipping of the ECG channel. For participants where QRS detection of the ECG channel failed, one method used in this study was to perform QRS detection on every EEG channel and select the channel containing the mode of the detected QRS complexes. From this channel, the median template is created and applied across channels for pulse artifact removal.

To address these numerous sources of noise, there are also techniques. Gradient artifacts can be minimized by modifying the configuration or layout of EEG leads^67^. To remove the gradient field artifact, we use the MR clock to record the scanner trigger at every TR^68,69^. Using a template artifact subtraction method^70^, the gradient artifact is recorded at each TR onset and averaged to create a template. The template is then subtracted from the signal to produce a clean signal. In this study, we used the MRIB plug-in for EEGLAB, provided by the University of Oxford Centre for Functional MRI of the Brain (FMRIB), to regress the MRI gradient artifact^36,37^. For noise induced by the helium compressor, there are methods for recording and regressing this motion induced artifact^7,71^; however, in our experiments, the simplest method for removing this artifact was to turn off the helium compressor during simultaneous recordings. While there is a risk of helium boiling off as temperature rises in the scanner, this can be addressed by having shorter scan sessions. In our study, the temperature of the cooling system did not fluctuate, which would impact cryogen loss. While shorter scans are ideal, we collected data upwards of 2 hours without issue.

Another factor found to impact EEG data quality was signal clipping, which often appears in the ECG channel during simultaneous recordings. In our simultaneous EEG-fMRI recordings, cable and amplifier placement inside the scanner affected EEG data quality. For cable placement, several factors must be taken into consideration. When scanning, researchers must ensure their setup minimizes loops, cables should run along the center of the bore, and the connected amplifier should be placed at the center of the bore to ensure better data quality^65^. Excessive bends or loops in wires can induce currents in the cables, thus introducing artifacts into the EEG signal. Another way to reduce artifacts is to reduce cable length between the EEG cap and connected amplifiers. All major scanner vendors offer head coils that are designed with a channel for EEG cables that lie directly above a participant’s head^72^. In addition, cables that are bundled produce less artifacts than ribboned cables^73^. In our experiments, the head coil did not contain a channel for EEG cables and a ribboned cable was used to connect the cap and amplifier. To reduce artifacts, EEG cables were run through the head coil above the participant’s head and taped along the center of the bore to minimize movement and to ensure optimal position in the scanner.

### Concluding remarks

Simultaneous EEG and fMRI offer rich data that provides complimentary temporal and spatial information. Here, we demonstrated the consistency of EEG results outside and inside the scanner. These analyses also indicate the utility of EEG-informed analyses that can provide fMRI that are data-driven. Additionally, the main benefit of this release is offering a rich resource of naturalistic viewing (Inscapes and movie clips), resting state data, and task data (checkerboard), that to our knowledge, is yet to be available. Investigators can also use this resource to optimize data processing pipelines and explore secondary analyses. Finally, this data release continues the trends of open data initiatives that provide transparency, efficiency, and an indispensable community resource.

## Usage Notes

Data for this study is shared using the Commons Attribution-ShareAlike 4.0 International (CC BY-SA 4.0) license. This license gives permission to copy and redistribute the data in any medium or format. It also gives permissions to remix, transform, and build upon the data for any purpose, even commercially. Anyone using this data must give appropriate credit, provide a link to the license, and indicate if changes were made. Additionally, any further distribution must be under the same license as the original dataset.

## Code Availability

Code for presenting task stimuli and naturalistic stimuli, along with code to preprocess EEG and fMRI imaging data, is available on GitHub at: https://github.com/NathanKlineInstitute/NATVIEW_EEGFMRI. Additionally, the videos used for naturalistic stimuli will also be made available through the GitHub repository.

## Acknowledgments

We would like to acknowledge Raj Sangoi and Caixia Hu for providing their technical support and expertise in developing the scanning protocol for data collection. We would also like to acknowledge Mark Higger for his contributions developing code for the EEG preprocessing pipeline.

